# Rapid molecular detection of Zika virus in urine using the recombinase polymerase amplification assay

**DOI:** 10.1101/078501

**Authors:** Ahmed Abd El Wahed, Sabri S. Sanabani, Oumar Faye, Rodrigo Pessôa, João Veras Patriota, Ricardo Rodrigues Giorgi, Pranav Patel, Susanne Böhlken-Fascher, Olfert Landt, Matthias Niedrig, de A. Zanotto Paolo M., Claus-Peter Czerny, Amadou A. Sall, Manfred Weidmann

## Abstract

**Background:** Currently the detection of Zika virus (ZIKV) in patient samples is done by real-time RT-PCR. Samples collected from rural area are sent to highly equipped laboratories for screening. A rapid point-of-care test is needed to detect the virus, especially at low resource settings.

**Methodology/Principal Findings:** In this report, we describe the development of a reverse transcription isothermal recombinase polymerase amplification (RT-RPA) assay for the identification of ZIKV. RT-RPA assay was portable, sensitive (21 RNA molecules), and rapid (3-15 minutes). No cross-reactivity was detected to other flaviviruses, alphaviruses and arboviruses. Compared to real-time RT-PCR, the diagnostic sensitivity was 92%, while the specificity was 100%.

**Conclusions/Significance:** The developed assay is a promising platform for rapid point of need detection of ZIKV in low resource settings and elsewhere (e.g. during mass gathering).

**Author Summary:** Currently, Dengue (DENV), Zika (ZIKV) and Chikungunya (CHIKV) viruses represent a global threat. The clinical picture of the acute febrile diseases caused by DENV, ZIKV and CHIKV is very similar, in addition, the same mosquito vector is involved in the transmission cycle. The differentiation between them is of great importance as supportive treatment differs and the identification of any of the three viruses prompts implementation of control measures to avoid spreading of an outbreak. We have developed an assay for the detection of ZIKV genome. The assay based on isothermal “recombinase polymerase amplification” assay, which was performed at one temperature (42°C). The result was obtained in maximum of 15 minutes. Moreover, the assay is easy to be implemented at low resource settings.

## Introduction

Zika virus (ZIKV) associated with severe congenital malformations (*e.g.*, microcephaly, hydrocephalus) and neuropathies (e.g. Guillian-Barré-like syndrome) has been declared a public health emergency of International concern by WHO [1, 2]. In the absence of a specific treatment and vaccine, early diagnostics is key to control the epidemic and trigger intervention. Diagnostic challenges are unspecific symptoms in the context of co-circulating Dengue virus and Chikungunya virus, limited laboratory infrastructure in rural areas where most cases occur, cross reactivity impeding serological detection and need for rapid detection. Here we report on the development and use of a simple, mobile, point of need molecular assay based on reverse transcription recombinase polymerase amplification (RT-RPA) assay for detection ZIKV in urine in 15 minutes.

## Methods

### Ethics statement

Human urine samples tested in Brazil were provided by the Laboratory of Medical Investigation in Dermatology and Immunodeficiency, LIM-56/03, São Paulo Institute of Tropical Medicine, Faculty of Medicine the University of São Paulo, São Paulo, Brazil, which has obtained the ethical approval from the Faculty of Medicine, São Paulo University Review Board (FMUSP N0 1.184.947). All study participants provided informed consent.

### Test development

In order to determine the analytical sensitivity of the RT-RPA assay, ZIKV NS1/NS2 molecular RNA standard was ordered at concentration of 10^10^/µl from GenExpress (Gesellschaft für Proteindesign mbH, Berlin, Germany). The RT-RPA primers and probe (FP: 5´-TCTCTTGGAGTGCTTGTGATTCTACTCATGGT-3´; RP, 5´-GCTTGGCCAGGTCACTCATTGAAAATCCTC-3´; exo-probe, 5´-CCAGCACTGCCATTGA(BHQ1-dT)(Tetrahydrofuran)(FAM-dT)GCTYATDATGATCTTTGTGGTCATTCTCTTC-phosphate-3´) were designed in NS2A region conserved among all ZIKV lineages (nt 3572 to 3713, GeneBank: LC002520.1). The European Network for Diagnostics of Imported Viral Diseases (ENIVD) provided flaviviruses, alphaviruses and arboviruses that were used in the testing the cross-reactivity of the ZIKV RT-RPA assay. The clinical performance of the assay was evaluated on acute-phase urine samples collected from suspected cases at the Municipal Hospital of Tuparetama, Pernambuco, Brazil. The nucleic acid extraction and RT-RPA assay were applied as previously described [3] and all results were compared with a real-time RT-PCR [4] as gold standard.

### Statistical methods

A semi-log regression analysis and a probit analysis were performed by plotting the RT-RPA threshold time (Tt) against the number of molecules detected to determine the ZIKV RT-RPA assay analytical sensitivity using PRISM (Graphpad Software Inc., San Diego, California) and STATISTICA (StatSoft, Hamburg, Germany), respectively. Diagnostic sensitivity and specificity were calculated using standard formulas. In addition, a linear regression analysis was performed using the values of real-time RT-PCR cycle threshold (Ct) and RT-RPA Tt by PRISM.

## Results

The limit of the detection of the assay was 21 RNA copies (95% probit analysis of dataset of eight RT-RPA runs using NS1/NS2 RNA diluted standard, Fig 1 and 2). The RT-RPA assay identified African (GenBank: AY632535) and Brazilian strains (Instituto Evandro Chagas, Belém, Brazil) down to 65 and 35 RNA genome equivalents, respectively, using ten-fold serial dilutions from virus culture supernatant.

**Fig 1.**
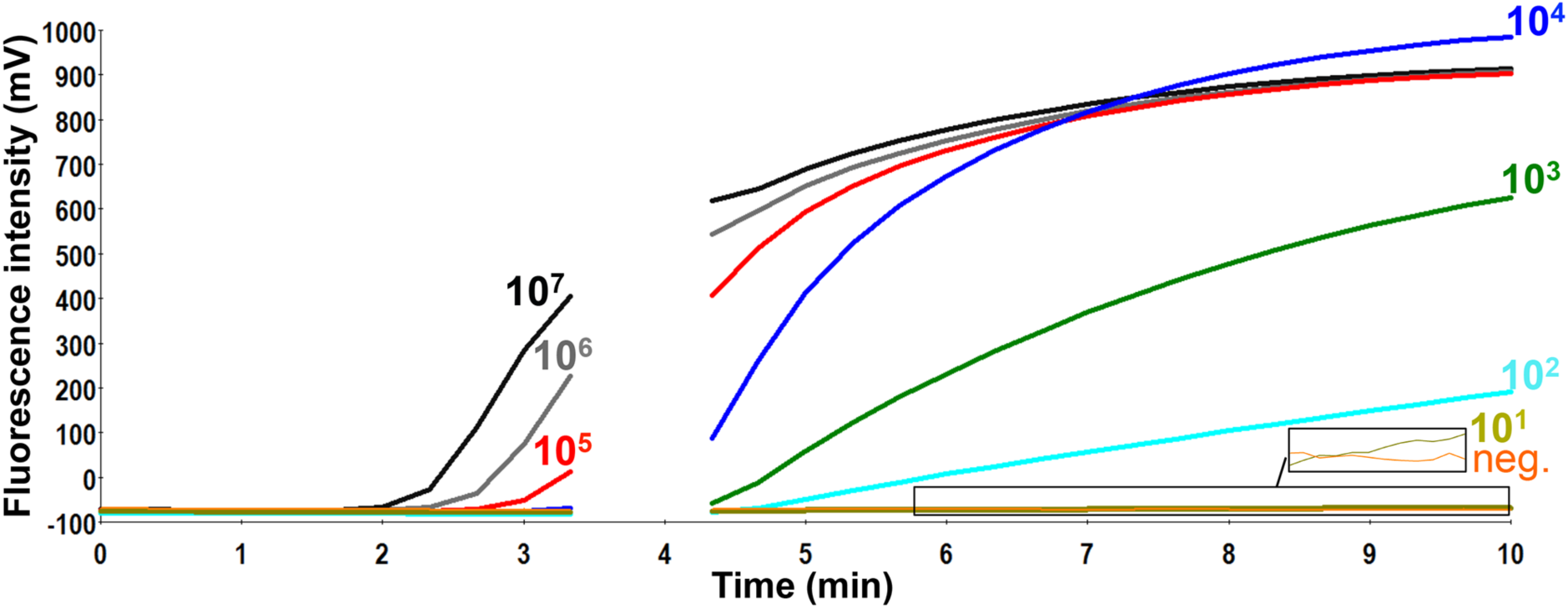
Analytical sensitivity of ZIKV RT-RPA assay. Fluorescence development *via* real-time detection in one RT-RPA run by using a dilution range of 10^7^-10^1^/µl of the RNA molecular standards (Graph generated by ESEquant tubescanner studio software). The limit of detection was 10 RNA copies. Data of eight RT-RPA runs is used for the probit regression analysis in Fig 2. The box in the lower right corner of the figure magnifies the fluorescence signals for the ten RNA copies and the negative control as the signal for ten RNA copies is very low.

**Fig 2.**
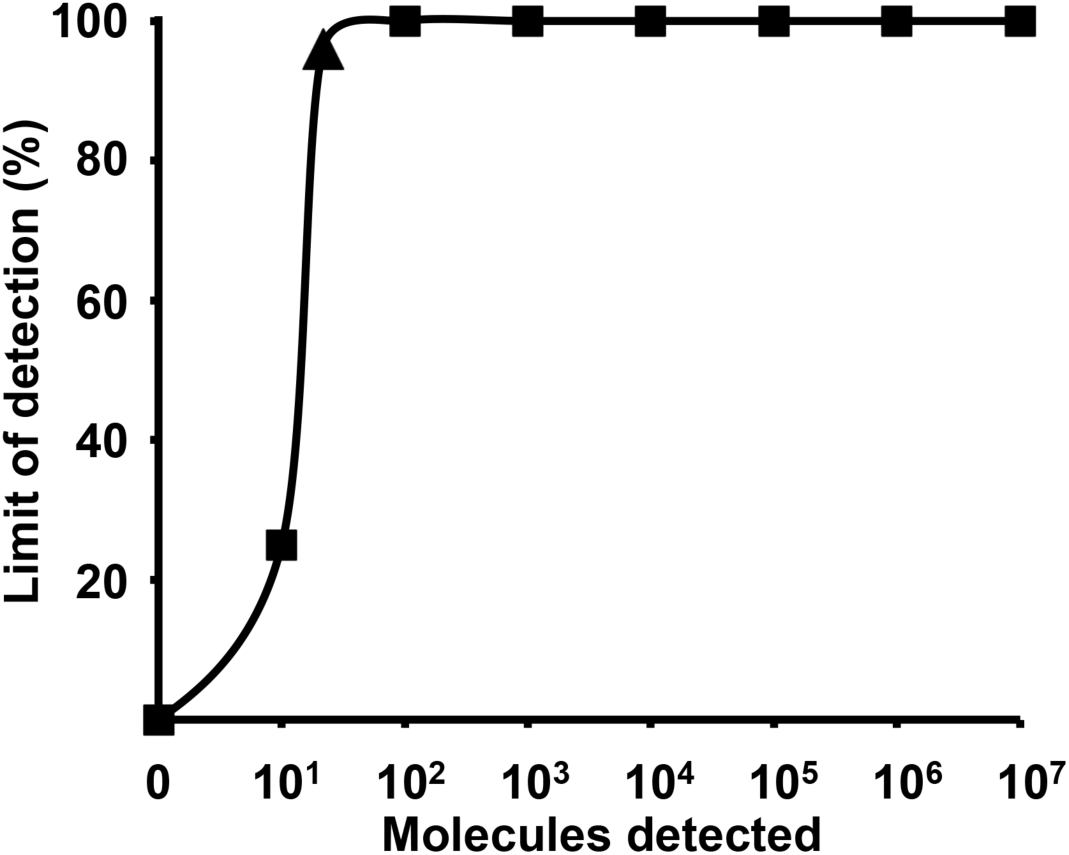
The probit regression analysis using data of eight RT-RPA assay runs. The limit of detection at 95% probability (21 RNA molecules) is depicted by a triangle. 10^7^−10^2^ RNA molecules were detected 8 out of 8 runs, while 10^1^ copies were identified 2 out of 8 runs of the ZIKV RT-RPA assay.

The assay is highly specific as no amplification was observed with the following viruses: Dengue 1-4, West Nile, Yellow Fever, Tick borne encephalitis, Japanese Encephalitis, Rift Valley Fever and Chikungunya. The clinical performance of the RT-RPA assay was tested using 25 positive (Ct values: 30–39, Fig 3) and nine negative urine samples collected during the ZIKV epidemic in Tuparetama, Brazil. The RT-RPA identified 23/25 (Sensitivity: 92%) positive (Fig 3) and 9/9 (specificity: 100%) negative samples.

**Fig 3.**
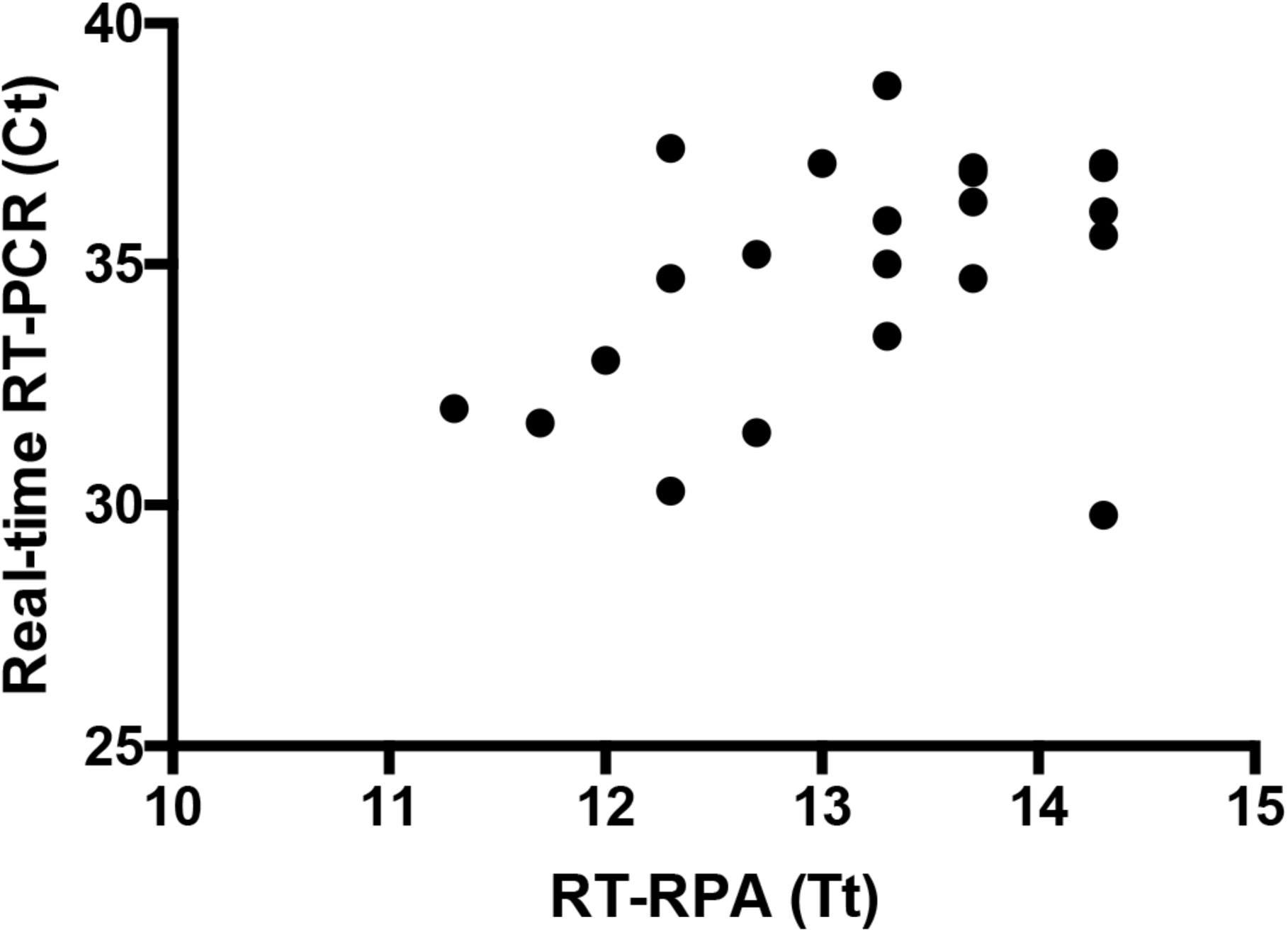
Results of screening 25 urine samples with both real-time RT-PCR and RT-RPA assays. Twenty samples are shown as three samples produced identical results and two were negative in RT-RPA assay.

## Conclusions

To our knowledge, the developed RT-RPA assay is the first sensitive rapid molecular assay applied on field samples for the detection of ZIKA in 15 minutes, which could be implemented at the point of need in a mobile suitcase laboratory [5] to make testing of pregnant women available directly in rural settings. Moreover, combining it with our developed dengue [6] and chikungunya [7] RT-RPA assays will allow its use during outbreak investigations.

### Box 1. Advantages and Disadvantages of the ZIKA RT-RPA assay

#### Advantages

1. Rapid (15 minutes) and limit of detection is 21 RNA copies
2. Highly specific
3. Easy to use at point of need as all reagents are cold chain independent, the assay is operated *via* a portable device and the cost per test is 4 Euro

#### Disadvantages

1. Obtaining sensitive RPA primer and probe is difficult as no strict rules for designing the RPA oligonucleotides are available
2. Slightly less sensitive than the real-time RT-PCR (diagnostic sensitivity is 92%)
3. Do not differentiate between the two lineages of the ZIKV

